# LZTS2 Emerges as a Regulator of Craniofacial Development and Modulator of DYRK1A

**DOI:** 10.64898/2026.03.31.715576

**Authors:** Nicole Cheng, Santiago Lima, Larisa Litovchick, Amanda JG Dickinson

## Abstract

**Background:** Precise control of DYRK1A dosage is essential for embryonic development, including craniofacial morphogenesis. While LZTS2 is among the most consistently identified DYRK1A-interacting proteins, its roles in embryonic development remain incompletely understood, and its potential contribution to craniofacial development has not been examined. *Xenopus laevis* was used to test the role of LZTS2 in craniofacial development and its functional relationship with DYRK1A.

**Results:** Lzts2 and Dyrk1a showed overlapping expression during craniofacial development, with both proteins present in developing facial tissues. Knockdown of Lzts2 disrupted craniofacial morphogenesis and reduced expression of the neural crest-associated genes *sox9* and *pax3*. These phenotypes closely resembled those caused by decreasing Dyrk1a function. Sub-phenotypic reductions of Lzts2 and Dyrk1a synergized to produce craniofacial defects, while partial reduction of Lzts2 attenuated aspects of the phenotype caused by Dyrk1a overexpression. Comparative analysis of human phenotypes associated with copy number gains of *LZTS2* and *DYRK1A* revealed striking overlap, consistent with a potential functional interaction between these genes in humans.

**Conclusions:** These findings identify Lzts2 as a previously unrecognized regulator of craniofacial development and support a functional interaction with Dyrk1a during embryogenesis. Modulating LZTS2 or related regulatory partners may provide a strategy to selectively tune DYRK1A-dependent developmental pathways

## INTRODUCTION

DYRK1A (Dual-specificity tyrosine-regulated kinase 1A) is a multifunctional and widely expressed protein in the embryo (Blackburn et al., 2019; Cho et al., 2019; Fotaki et al., 2002; Johnson et al., 2024; Willsey et al., 2020). As a central signaling regulator present across tissues and developmental stages, DYRK1A sits at the crossroads of proliferation, differentiation, and morphogenesis. It is therefore unsurprising that humans are exquisitely sensitive to changes in Dyrk1a dosage (Ananthapadmanabhan et al., 2023; Atas-Ozcan et al., 2021; Dierssen and de Lagrán, 2006; Duchon and Herault, 2016). Indeed, gene copy number variations and *de novo* mutations in DYRK1A cause DYRK1A Haploinsufficiency Syndrome, a disorder affecting the facial region, brain, and multiple organ systems (Becker et al., 2014; Ji et al., 2015; van Bon et al., 2016; Widowati et al., 2018). Conversely, increased dosage of DYRK1A, due to its location within the critical region of chromosome 21, is thought to contribute substantially to key manifestations of Down Syndrome, including craniofacial differences and intellectual disability (Antonarakis et al., 2020; Atas-Ozcan et al., 2021; Duchon and Herault, 2016; Kaczorowska et al., 2019; McElyea et al., 2016; Redhead et al., 2023). Beyond developmental disorders, DYRK1A has emerged as a significant player in cancer and other diseases (Fernandez-Martinez et al., 2015; Kim et al., 2021). Together, these findings underscore a central principle: DYRK1A dosage must be precisely controlled and understanding how its activity is regulated is critical for both developmental biology and disease therapeutics.

Recently, we and others have identified novel DYRK1A binding partners that may serve as key regulatory nodes (Ananthapadmanabhan et al., 2023; Guard et al., 2019; Menon et al., 2019a; Roewenstrunk et al., 2019). One of the most consistently reported binding partners of DYRK1A is LZTS2 (leucine zipper tumor suppressor 2). Examination of curated protein interaction datasets in the BioGRID revealed that LZTS2 ranks among the top DYRK1A-associated proteins, representing the third most frequently reported interaction partner (Oughtred et al., 2021). Across the database, a total of 12 independent experimental observations support an interaction between LZTS2 and DYRK1A. These include multiple affinity capture–mass spectrometry datasets as well as orthogonal validation approaches such as affinity capture followed by western blotting, fluorescence resonance energy transfer (FRET), and proximity labeling–mass spectrometry (Golkowski et al., 2023; Huttlin et al., 2021; Huttlin et al., 2017; Huttlin et al., 2015; Li et al., 2015; Menon et al., 2019b; Varjosalo et al., 2013; Wang et al., 2024; Youn et al., 2018). Notably, the interaction has been detected repeatedly across independent studies and experimental platforms, providing strong evidence that LZTS2 and DYRK1A physically associate in cells.

LZTS2 is a member of the Fezzin protein family (Wendholt et al., 2006), characterized by a conserved Fez1 DNA-binding domain and coiled-coil regions that enable homo- and heteromultimerization, as well as interactions with Spine-associated Rap GTPase-activating proteins (SPARs) (Dolnik et al., 2016; Spilker and Kreutz, 2010). These structural features position LZTS2 as a potential scaffold and signaling integrator. Functionally, LZTS2 has been implicated in diverse cellular processes, including regulation of β-catenin signaling, axon outgrowth, microtubule severing, central spindle formation, and cytokinesis (Li et al., 2011; Liu et al., 2024; Sudo and Maru, 2008; Thyssen et al., 2006; Yu et al., 2017). Consistent with these roles in cell division and signaling, LZTS2 deregulation is associated with multiple cancers (Arnold et al., 2006; Johnson et al., 2013; Tang et al., 2024; Yu et al., 2017). Importantly, LZTS2 is expressed during embryogenesis and has been implicated in aspects of vertebrate development, yet its developmental functions remain incompletely defined due to the limited number of studies in embryonic systems (Gessert et al., 2011; Li et al., 2011; Peng et al., 2011; Yu et al., 2017).

DYRK1A is well established as a regulator of craniofacial development (Johnson et al., 2025; Johnson et al., 2024; McElyea et al., 2016; Redhead et al., 2023), yet despite LZTS2 being a known binding partner, its role in craniofacial morphogenesis remains unexplored, and whether these proteins functionally interact during this process is unknown. To address this, we investigated the role of LZTS2 in craniofacial development in *Xenopus laevis* and examined its potential to influence DYRK1A function during embryogenesis.

## MATERIALS AND METHODS

### Obtaining *Xenopus laevis* embryos

*Xenopus laevis* embryos were obtained using standard procedures (Sive et al., 2000) approved by the VCU Institutional Animal Care and Use Committee (IACUC protocol number AD20261). Embryos were staged according to Nieuwkoop and Faber (Nieuwkoop and Faber, 1994). Stages are also reported as hours post fertilization at 23C.All studies are performed on stages from fertilization to stage 42-43 (∼80-85hpf at 23C). By stage 43, the main vesicles of the brain are apparent, eyes have a retina and lens, the heart is beating and blood is circulating, much of the cranial cartilage has formed and the digestive system is still taking shape. We generally refer to the stages used in this range (up to stage 43) as embryos because they are similar to and undergoing the same processes as human embryos.

### Morpholino Knockdown of Lzts2 in *Xenopus laevis*

To knockdown gene function we used antisense oligonucleotides stabilized with morpholino rings (Morpholinos (MOs) GeneTools). Since *Xenopus laevis* is tetraploid and thus has two homeologs of each gene (denoted S and L) we designed two Lzts2 morpholinos (MOs) that targeted splice junctions

(LZTS2.LMO; ATCATGTTATTCTCGTTTACCTGGC, LZTS2.SMO; TAACAAACATCCTGCCGTACCTGG).

We also used a previously validated translation blocking Dyrk1 MO that target both homeologs of this gene (Xenbase.org (Blackburn et al., 2019; Johnson et al., 2024)). A standard control MO that does not target any *Xenopus laevis* sequence was utilized as a negative control. MOs were diluted to 34ng/ul to make stock solutions and from there further diluted to achieve the concentrations indicated in the results. 1-5nl were injected into each embryo. All MOs were labeled with fluorescein which allowed separation of un-injected individuals by 24 hours of development. Successfully injected embryos, as indicated by green fluorescence and being alive, were sorted under a stereoscope and raised at 15C in frog embryo media (0.1XMBS). To assess whether the *lzts2* MOs resulted in changes in mRNA sequence, PCR was performed with Apex Hotstart Taq master mix (Bioline, cat # 42-144) on a BioRad MJ Mini Personal Thermocycler. The PCR products were analyzed on a 1% agarose gel prepared with molecular grade agarose (Bioline, cat # BIO-41025) in TAE buffer. Primers targeting the flanking exons were used to detect splicing defects (*lzts2*.*L* left; TACCAATCAGCCCCTCTAGC, right TCCATTTCCTGCTCACACCT and *lzts2*.*S* left; ATCAGCCCCTCTAGTGATGC, right; CTCCTCCATTTCTTGCTCGC).

### Crispr/Cas9 mutagenesis of *lzts2* in *Xenopus laevis*

sgRNA targeting each homeolog of lzts2 was designed using the ChopChop software. Sequences were chosen that best targeted the desired gene with no off-targets and high efficiency (target sequences = lzts2.S; TATTCTGATCGAGCATCCAGGGG, lzts2.L; CATCGTTCAAGATCGGGGAGCGG). The sgRNAs were purchased from Synthego and diluted as recommended in low EDTA TE buffer. Then, 200 pg of sgRNA was incubated with 2 pg of Cas9 protein (PNA Bio Inc., cat # CP01) for 10 min and then 1-2ul was injected into each embryo. A negative control consisted of the same concentrations of Cas9 protein. Since there was no way to visually determine whether an embryo was mutant, all embryos injected were counted but only those with some types of malformation were used to quantify defects. To confirm mutations, the DNA was extracted from 8-10 randomly selected embryos (with craniofacial malformations) using the HotShot protocol. Each embryo was immersed in 40 μl of an alkaline lysis buffer (25 mM NaOH, 0.2 mM Na-EDTA) and heated for 40 min at 95°C. The solution was then cooled and an equal volume of neutralization buffer (40 mM Tris–HCL) was added. One microliter of this solution was used in a standard PCR reaction with Hotstart Taq master mix and primers that flanked the predicted mutation site. The product was then sent for purification and sequencing at Genewiz (Azenta Life Sciences, South Plainfield, NJ).

### dyrk1a mRNA synthesis for injection in Xenopus laevis

Plasmids containing the *Xenopus laevis dyrk1a* containing plasmid was a kind gift from Toshiyasu Goto and GFP containing plasmid from Dr. Hazel Sive. The plasmids were both linearized with NotI, and mRNA synthesized using the SP6 mMessage Kit following the recommended directions.

### Microinjection and *Xenopus laevis* embryo culture

Microinjection of reagents were performed using an Eppendorf Femtojet micro-injector and a Zeiss Discovery V8 stereoscope. Embryos were placed in a dish lined with nylon Spectra mesh (1,000 um opening and 1350 um thickness) at the bottom to hold embryos in place and filled with 3% Ficoll 400 (Fisher, cat # BP52 5) dissolved in frog embryo media (0.1X MBS (Modified Barth’s Saline)). After injections embryos were randomly divided into 3 groups and placed into 10cm culture dishes filled with frog embryo media and maintained in 15C incubator. Media was changed daily, and dead embryos removed.

### qRT-PCR in *Xenopus laevis*

Expression of *sox9, pax3* and *dyrk1a* was assessed using quantitative reverse transcription PCR (RT-qPCR). RNA was extracted from the heads of embryos at stage 32 using TRIzol extraction followed by lithium chloride precipitation. One-step quantitative RT-PCR was performed using the Luna One-Step Universal RT-qPCR kit (NEB, E3005S) and carried out on the Mic qPCR Machine (BioMolecular Systems) using the primer sequences previously published(Johnson et al., 2024). Using the delta-delta CT method, the relative fold change between control and experimental groups in the PCR products was calculated using *odc* as a reference gene.

### Immunofluorescence in *Xenopus laevis*

To perform all fluorescent labeling, embryos were fixed in 4% paraformaldehyde. To create transverse sections through the oral cavity, fixed embryos were immersed in 5% low melt agarose, and 200micron sections created with a vibratome (Leica). The sections were incubated in blocking buffer (1% Triton-X, 1% normal calf serum, 1% BSA in PBT) overnight at 4C. This was followed by primary antibody incubation (anti-DYRK1A and anit-LZTS2; Bethyl Laboratories, 1:100) for 48 hours. Sections were then incubated in secondary antibody (1:500, goat-anti mouse Alexa Fluor 488 or 568) combined with 2 drops of NucBlue (Invitrogen, RF37606), for 24-48 hours. Fluorescent images were collected with a C2 Nikon confocal microscope using 0.5-1micron steps and compiled using the maximum intensity function to compress the z stacks in the Nikon Elements software.

### DYRK1A Pharmacological antagonist treatments in *Xenopus laevis*

Embryos were immersed in 25uM INDY with 0.1% DMSO in 4 ml of frog embryo media (0.1X MBS) as described previously (Johnson et al., 2024). Controls were immersed in 0.1% DMSO alone in the same volume. Embryos were treated at stage 22-24 (∼24hpf) for 24hrs at 23C and then washed out and cultured until the desired stage in frog embryo media.

### Imaging *Xenopus laevis*

At stage 43, tadpoles were anesthetized in 1% tricaine for 10 minutes and then fixed in 4% paraformaldehyde overnight at 4°C. Whole embryos were imaged and the heads were removed to effectively view and image the face and head. A No. 15 scalpel (VWR, cat. no: 82029-856) and Dumont No. 5 forceps (Fisher, cat. no: NC9404145) were used to make two cuts to isolate the head: first-at the posterior end of the gut and then second caudal to the gills. Isolated heads were mounted in small holes or depressions in either agarose or clay-lined dishes containing Phosphate Buffered Saline with 0.1% Tween (PBT). The faces were imaged using a Discovery V8 stereoscope fitted with an Axiovision digital camera (Zeiss).

### Measurements and Statistics in *Xenopus laevis*

Measurements of intercanthal distance, the distance from the inside of each eye across the midface was measured using Zeiss Zen software. The mouth roundness was measured in ImageJ using the “roundness” function where Roundness = 4*Area/(pi*MajorAxis^2). Data was imported into excel to create stacked bar graphs. The entire range of measurements for each dataset (including controls and experimental) was divided evenly into three groups (small, medium, large). 60 embryos in 2-3 biological replicates were quantified for each treatment group. SigmaPlot was utilized to perform statistical analyses and to create box and whiskers plots. A multiple group analysis was performed where it was first analyzed for normality and equal variance. An ANOVA or ANOVA on Ranks (Kruskal Wallis test) were performed to determine if there were statistical differences across all treatment groups. This was then followed by a Tukey posthoc test to compare two groups. Significance was deemed if p values were lower than 0.05.

### Plotting *Xenopus laevis* protein and mRNA levels over time in Xenbase

Xenbase.org was used to plot relative Dyrk1a protein and dyrk1a mRNA expression. This function generates the plot for a desired gene based on published proteomic and transcriptome data (Fisher et al., 2023).

### Analysis of *Xenopus laevis* Lzts2 DNA and protein sequence and structure

Xenopus laevis lzts2 sequences were retrieved from Xenbase.org (Fisher et al., 2023). The mRNA sequences of lzts2.L (XM_041568699.1) and lzts2.S (XM_041570954.1) were aligned using Multalign (http://multalin.toulouse.inra.fr/multalin/). Lzts2.L protein (XP_018080722.1) was similarly aligned with LZTS2 human protein (AAH58938.1). Protein domains created from information at NCBI. 3D models of X. laevis and human Lzts2 proteins were predicted using default settings of Swiss-Model (https://swissmodel.expasy.org)

### DECIPHER analysis of Genotype-Phenotype data

DECIPHER (DatabasE of genomiC varIation and Phenotype in Humans using Ensembl Resources)(Firth et al., 2009) was used to collate phenotypes in two or more patients with DYRK1A and LZTS2 copy number gains. Phenotype comparison was performed using compare lists program by the Bioinformatics and Computing (BARC) tool developed and hosted by the Whitehead Institute (Whitehead BaRC public tools (mit.edu)),

## RESULTS

### 1. LZTS2 is expressed during craniofacial development and overlaps with Dyrk1a expression

*Xenopus laevis* is both an effective model to study craniofacial development as well as the ideal vertebrate model to test for protein interactions *in vivo*. If Lzts2 interacts with Dyrk1a in Xenopus embryos then we would expect that these proteins would be present at the same time and in the same tissues during development. The temporal expression of both mRNA and protein levels of Dyrk1a and Lzts2 were analyzed using both proteomics and RNA-seq data available at Xenbase.org. Results indicated that mRNA and protein expression of *lzts2*/Lzts2 and *dyrk1a*/Dyrk1a followed very similar temporal profiles over time (Fig.1Ai-ii). In addition, the reported expression of *lzts2* mRNA appears similar to *dyrk1a* in *Xenopus* embryos, notably in the branchial arches (Blackburn et al., 2019; Gessert et al., 2011). To extend this localization data we also compared Dyrk1a and Lzts2 protein expression in the developing head using mammalian antibodies that recognize conserved regions of the proteins. Specificity of the antibodies were determined in knockdown experiments (see below and (Johnson et al., 2024)). Our results revealed widespread expression of both proteins throughout the face including the epidermis, cranial muscles and mesenchymal cells of the early tadpole stages when facial tissues are developing (Fig. 1B). In the mesenchymal cells, derived from neural crest, Dyrk1a was strongly expressed in nuclei and especially cells appearing to be undergoing cell division (Fig. 1B, pink arrow). But both Lzts2 and Dyrk1a appear in the cytoplasm of these mesenchymal cells (Fig. 1B, white arrows). Lzts2 and Dyrk1a were also both localized to multiciliated cells in the epidermis (Fig. 1B, white arrowheads). In some tissues Lzts2 and Dyrk1a were enriched in different cellular compartments. For example, in the cranial muscles Dyrk1a was strongly expressed in the nuclei and less so in the cytoplasm whereas Lzts2 appears more strongly expressed in the cytoplasm.

**Figure 1.**
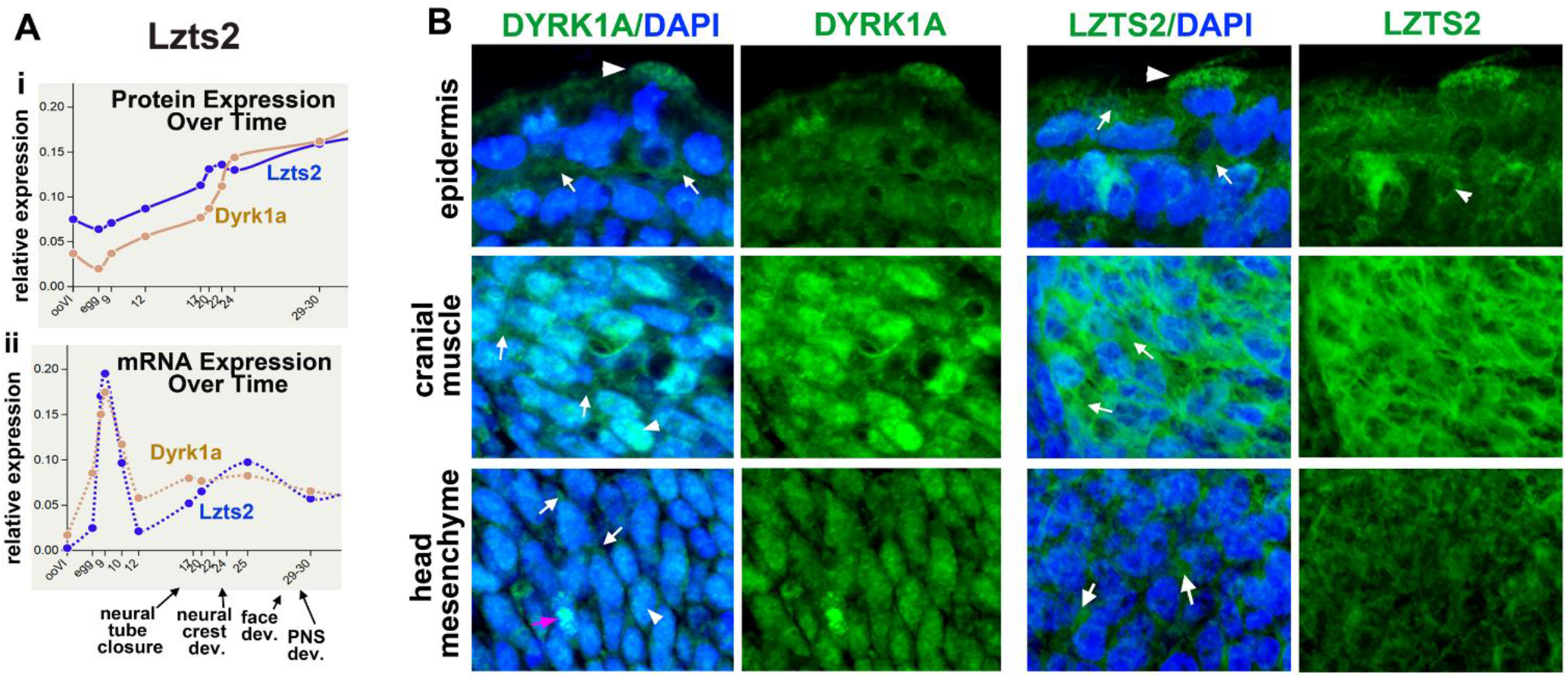
Lzts2 and Dyrk1a show overlapping temporal expression and localization in craniofacial tissues. **A***) Temporal expression of Lzts2 and Dyrk1a during embryonic development*. (i) Relative protein expression levels of Lzts2 and Dyrk1a across developmental stages generated from Xenbase proteomics datasets. (ii) Relative mRNA expression profiles of lzts2 and dyrk1a across embryonic development derived from Xenbase RNA-seq data. Key developmental transitions including neural tube closure, neural crest development, facial development, and peripheral nervous system (PNS) development are indicated. **B)** *Localization of Dyrk1a and Lzts2 in craniofacial tissues*. Immunofluorescence labeling of Dyrk1a and Lzts2 in transverse sections of the head from stage 40 embryos (≈60 hpf). Images show merged channels with DAPI nuclear counterstain (blue) and antibody labeling (green), as well as antibody signal alone. Dyrk1a and Lzts2 are detected in multiple craniofacial tissues including the epidermis, cranial muscle, and cranial neural crest–derived head mesenchyme. White arrows indicate cytoplasmic labeling, arrowheads indicate nuclear localization, and the pink arrow indicates a mitotic cell exhibiting strong nuclear Dyrk1a signal.

Together the overlapping temporal and spatial expression of Dyrk1a and Lzts2 supports a possible connected function in craniofacial tissues. Future work and compatible antibodies are required to perform cololocalization studies to pinpoint more precisly when and where Lzts2 and Dyrk1a are co-localized in craniofacial cells.

### 2. Lzts2 knockdown is specific and causes developmental differences

To determine whether Lzts2 is required for craniofacial development we used antisense oligos with a stabilizing morpholino ring; Morpholinos (MOs) to decrease the functional protein in the embryo. The two *lzts2* homeologs (*lzts2*.*L* and *lzts2*.*S*) are 91% identical (Suppl Fig 1A). In addition, the proteins encoded by each *lzts2* homeolog are expressed at almost identical levels over early embryonic development suggesting shared, overlapping and/or redundant functions (Suppl Fig. 1B). Regardless, two morpholinos were designed to bind to splice junctions of each of the *lzts2* homeologs. MOs were labeled with fluorescein to track the injections and allow the removal of any uninjected embryos. A standard control fluorescein labeled MO (40-80ng) or both *lzts2*.*S* and *lzts2*.*L* MOs combined (20-40ng of each) were injected into embryos at the 1 cell stage (Fig 2A). At stage 22 (∼24hrs @23C), such embryos with green fluorescence were used to validate the effectiveness of the MOs. Since our *lzts2* MOs were designed to target splice junctions, they were predicted to result in mis-splicing which would result in changes in transcript size. To confirm this prediction, RT-PCR was performed using primers that bind to exons flanking the MO target site. Results indicated that indeed smaller PCR products were identified in the *lzts2* morphants consistent with mis-splicing (Fig.2A pink asterisks). Mis-spliced gene products can become degraded and therefore these morpholinos could reduce the Lzts2 protein. To determine if the proteins were indeed decreased, we assessed their levels by immunofluorescence using a human antibody predicted to bind to both Lzts2 homeologs. Fluorescein labeled Lzts2 MOs were injected into one cell at the 2 or 4-cell stage to create partial or mosaic *lstz2* morphants. After 24 hours (@23C) the embryos were fixed, sectioned, and labeled. Results indicated that LZTS2 antibody labeling (pink) appeared to be decreased in cells positive for the morpholino (green) (Fig. 2B). Together these results indicate that *lzts2* morpholinos effectively reduced Lzts2 protein in the developing embryo.

**Figure 2.**
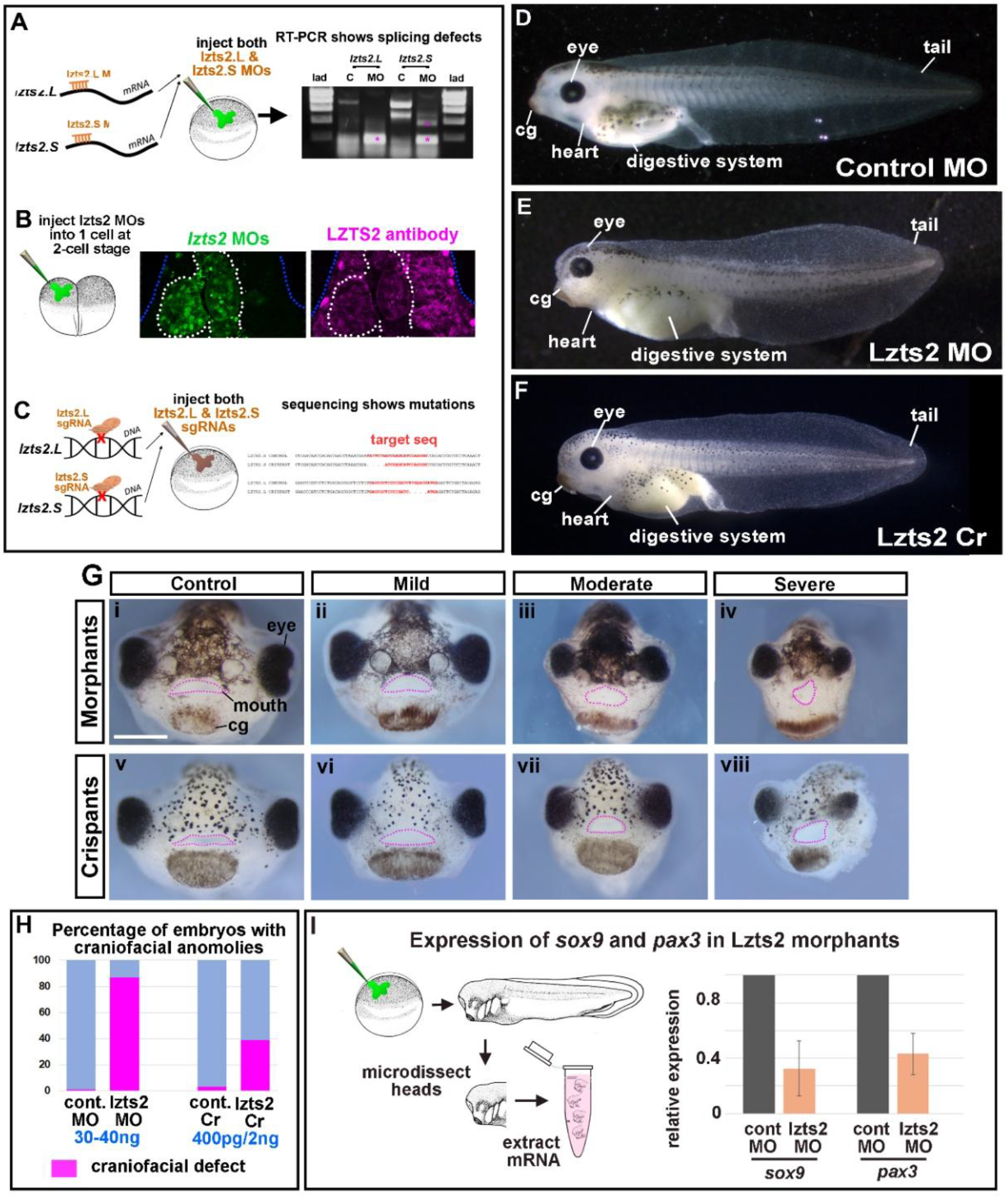
Lzts2 knockdown causes developmental and craniofacial anomalies in Xenopus embryos. **A)** *Morpholino-mediated knockdown of Lzts2 disrupts transcript splicing*. Schematic showing splice-blocking morpholino (MO) target sites in the lzts2.L and lzts2.S transcripts and injection into one-cell stage embryos. RT-PCR analysis demonstrates altered PCR products in Lzts2 morphants compared with controls, indicating mis-splicing of lzts2 transcripts. Asterisks mark abnormal PCR bands consistent with mis-spliced products. **B)** *Lzts2 protein is reduced in cells containing Lzts2 morpholino*s.Fluorescein-tagged Lzts2 MOs were injected into one blastomere at the two-cell stage to generate mosaic embryos. Immunofluorescence labeling using an LZTS2 antibody (magenta) shows reduced Lzts2 protein in MO-positive cells (green), confirming effective knockdown of the protein. **C)** *CRISPR/Cas9 mutagenesis of lzts2*. Schematic showing sgRNA target sites within the lzts2.L and lzts2.S genes and injection of sgRNAs together with Cas9 protein into one-cell stage embryos. Sequencing of PCR-amplified genomic regions confirms mutations at the targeted loci. **D–F)** *Lzts2 depletion causes developmental abnormalities*. Lateral views of representative embryos injected with control MO (D), Lzts2 MOs (E), or Lzts2 sgRNA/Cas9 reagents (F). Lzts2-deficient embryos exhibit developmental abnormalities affecting multiple tissues including the heart and digestive system. Abbreviations: cg, cement gland. (G) *Lzts2 depletion causes craniofacial defects*. Frontal views of representative embryos. (i) Control embryo. (ii–iv) Lzts2 morphants displaying increasing severity of craniofacial defects categorized as mild, moderate, and severe. (v) Cas9 control embryo. (vi–viii) Lzts2 crispants showing comparable craniofacial abnormalities. Dashed lines indicate the mouth outline. (H) *Quantification of craniofacial anomalies*. Percentage of embryos displaying craniofacial defects following Lzts2 MO injection (30–40 ng) or Lzts2 CRISPR/Cas9 mutagenesis. (I) *Neural crest marker expression is reduced in Lzts2 morphants*. Experimental schematic illustrating microdissection of embryo heads followed by RNA extraction and RT-qPCR analysis. Relative expression of the neural crest markers sox9 and pax3 is reduced in Lzts2 morphants compared with control embryos. Abbreviations: cg, cement gland; C, control; MO, morpholino; seq, sequence.

To further validate the specificity of the MO knockdown we also used an alternate knockdown method, Crispr/Cas9 mosaic mutagenesis in the F0 generation. 200 pg of each of the *lzts2*.*S* and *lzts2*.*L* gRNAs were combined (or a negative control sgRNA) and were injected with 2ng of Cas9 protein at the one-cell stage (Fig. 3C). A subset of the embryos with malformations were utilized for sequencing to confirm the Crispr/Cas9 induced mutation. To do so we used primers that targeted a 300-400 bp region surrounding the predicted mutation site in either *lzts2*.*L* or *lzts2*.*S*. Results indicated that as expected the PCR products had changes in the sequences at the sgRNA targets sites (representative sequences and mutations in Fig 2C).

**Figure 3.**
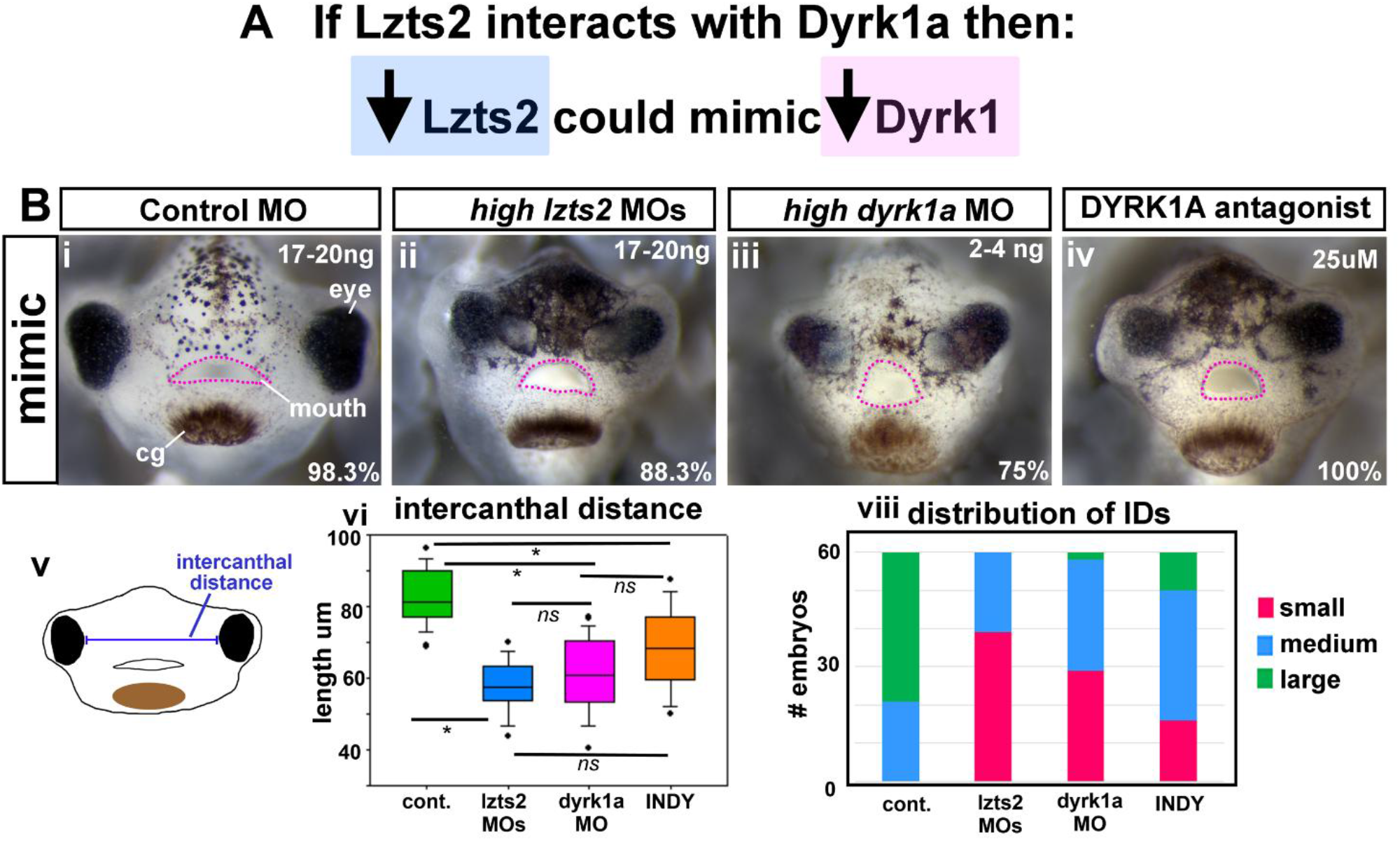
A) Lzts2 depletion phenocopies reduced Dyrk1a. **B)** (i–iv) Frontal views of representative embryos injected with control morpholinos (MOs), high-dose lzts2 MOs, high-dose dyrk1a MOs, or embryos treated with the DYRK1A inhibitor INDY. Dashed lines indicate the outline of the mouth. (v) Schematic illustrating measurement of the intercanthal distance (distance between the eyes). (vi) Quantification of intercanthal distances in control embryos and embryos with reduced Lzts2 or Dyrk1a activity (n = 60 embryos per condition). (viii) Distribution analysis of intercanthal distances displayed as stacked bar graphs, where values are divided into small, medium, and large categories.

We then compared the whole embryo phenotypes of the *lzts2* morphants and crispants. Lzts2 morpholinos were injected at three different concentration ranges that resulted in mild (15-20ng total), moderate (30-40ng total) and severe defects (60-80ng total). At the moderate range embryos were shortened and had digestive tract and heart anomalies as well as a smaller head with a more flattened face (Fig. 2D,E). Lzts2 Crisper/Cas9 reagents, injected as one standard concentration mix, caused mosaicism in the F0 generation and therefore produced embryos that range in severity. Lzts2 crispants that had a moderate phenotype appeared similar to moderate *lzts2* morphants (Fig. 2D-F). These results provide evidence that our knockdown methods are specific. Further, we also demonstrate that Lzts2 is required for the development of multiple systems consistent with what has been reported in other animal models.

### 3. Lzts2 knockdown causes craniofacial differences

L*zts2* mRNA and protein are present in craniofacial cells, despite this no studies to date have characterized craniofacial development in LZTS2 knockouts/knockdowns. Therefore, we next assessed craniofacial morphology in the mild, moderate and severe *lsts2* morphants and crispants. For this analysis we examined stage 43 when the mouth and jaws have formed. On average 99% of control embryos appeared normal. The 1% of abnormal control embryos were smaller but had no change in the craniofacial structures. 32% of embryos injected with a total of 15-20 ng of *lzts2* MOs had mild craniofacial malformations including an abnormally shaped mouth and eyes (Fig. 2Gi,ii and H). Doubling the MO concentration (total 30-40ng) resulted in 87% of embryos with craniofacial defects that included a narrower, rounder mouth, and smaller eyes (Fig. 2Giii and H). Finally, when injected with 60-80ng the embryos had similar malformations but were more severe (Fig. 2Giv, H). Interestingly, a subset of these severe embryos had asymmetrical malformations in the shape of the mouth, and one eye was more severely affected. We next compared *lzts2* Morphants with *lzts2* Crispants. We determined that 39% of embryos injected with a combination of *lzts2*.*S* and *lzts2*.*L* gRNAs (400pg total) with Cas9 protein had craniofacial defects, this is consistent with mutagenesis rates in our previous work (Bharathan and Dickinson, 2019). Such malformations ranged from mild to severe, and craniofacial defects included a narrower midface and rounder mouth (Fig. 2Gv-viii, and H). Importantly, many of the *lzts2* crispants closely resembled the *lzts2* morphants. No craniofacial abnormalities were noted in the negative control injected embryos except that some (∼1%) that were overall smaller in size (Fig. 2Gv, H).

Cranial neural crest cells are critical for giving the face its form and function. Sox9 and Pax3 transcription factors are important for neural crest development and were reduced in embryos treated with a Dyrk1a antagonist (Johnson et al., 2024). Therefore, we also examined the expression of these genes in *lzts2* morphants. Results indicated that indeed *sox9* and *pax3* expression levels were reduced by 0.33 and 0.42 respectively in embryos injected with *lzts2* MOs (Fig. 2I).

In summary, the results presented here demonstrate that indeed Lzts2 is required for craniofacial morphology and neural crest gene expression.

### 4. Lzts2 knockdown causes craniofacial phenotypes that mimic a reduction in Dyrk1a

If Lzts2 is required for Dyrk1a function in the embryo we predicted that a decrease in Lzts2 would cause some of the same developmental malformations as Dyrk1a inhibition or morpholino knockdown (Fig. 3A). To test this, we compared the *lzts2* morphants with embryos injected with *dyrk1* MO (8.5ng/embryo) or exposed to a Dyrk1a antagonist (INDY, 25uM). Remarkably, a subset of these embryos had very similar craniofacial appearance (Fig. 3Bi-iv). In particular, we noted similar narrower faces with closer set often angled eyes and mis-shaped mouths. To quantify this observation, we assessed intercanthal distance (distance between the eyes) that reflects the narrowing of the midface (Fig. 3Bv). The average intercanthal distance of *lzts2* morphants was 30.4% less than the sibling controls which was significant (p<0.001; Fig. 3Bvi). Dyrk1a morphants had a similar significant reduction of 26.0% (p<0.001; Fig. 3Bvi) and embryos treated with INDY had a significant 17.1% reduction in the average intercanthal distance (<0.001; Fig. 3Bvi). Further, the intercanthal distances of *lzts2* morphants were not different than the *dyrk1a* morphants (p=0.249; Fig. 3Bvi). We did find that there was a statistical difference in the average intercanthal distances of embryos treated with INDY and *lzts2* morphants (p<0.001; Fig. 3Avi). These results may reflect the more moderate phenotype we observed at the concentration of INDY used to treat embryos. A distribution analysis of intercanthal distances was performed where they were divided equally into three groups, small, medium and large (Fig. 3Bvii). Normally control embryos can vary in intercanthal distance and fall into both medium and large categories (representative image of a “large” intercanthal distance in a control embryo shown in Fig. 3Bi). However, this distribution shifted to include small intercanthal distances in *lzts2* morphants, *dyrk1a* morphants and INDY treated embryos (representative images of “small” intercanthal distances in Fig. 3Bii-iv and vii). Embryos exposed to INDY did have more embryos that fell into the “medium” category consistent with our qualitative observations (Fig. 3Bvii).

In summary, embryos with decreased *lzts2* or *dyrk1a* had strikingly similar facial malformations in the shape of the eyes and mouth as well as the narrowing of the midface. These data suggest that Lzts2 and Dyrk1a could have overlapping roles in craniofacial development.

### 5. Sub-phenotypic concentrations of dryk1a and lzts2 MOs synergize to cause craniofacial defects

If Lzts2 modulates Dyrk1a during development, then we hypothesized that knockdown of both proteins at the same time would synergize and cause malformations (Fig. 4A). To test this hypothesis a sensitized phenotypic assay was performed focusing on craniofacial morphology. To do this assay, low sub-phenotypic concentrations of *lzts2* and *dyrk1a* MOs were injected in combination. We predicted that if the two proteins work together to modulate the same process or pathway, we would observe a synergistic effect (Wahl et al., 2015). When embryos were injected with low concentrations of *lzts2* MOs (8-10 ng/embryo) or *dyrk1a* MO (2.5ng/embryo), no difference in craniofacial morphology was observed when compared to controls (compare Fig. 4Bi to Fig. 4Bii and iii). This was reflected in the insignificant minor differences in the intercanthal distance between the control and the low *lzts2* and (2.9%, p=0.465; Fig 4Bvi) and low *dyrk1a* (1.9%, p=0.793; Fig. 3Bvi). However, when *lzts2* and *dyrk1a* MOs were combined, 40% of these double morphants had clear qualitatively observable craniofacial defects with smaller faces and abnormal eyes as well as mis-shaped mouths (Fig. 4Biv). Quantification of intercanthal distances showed that there were significant differences between the double morphants compared to single *dyrk1a* morphants (17.0% decrease, p<0.001; Fig. 4Bvi) or single *lzts2* morphants (17.0% decrease, p<0.001; Fig. 4Bvi). These results were also evident when the distributions of intercanthal distances were plotted as stacked bar graphs (Fig. Fig. 4Bvii). In the double morphants there was a shift to include more embryos with “small” intercanthal distances compared to the single (*dyrk1a* or *lzts2*) morphants.

**Figure 4.**
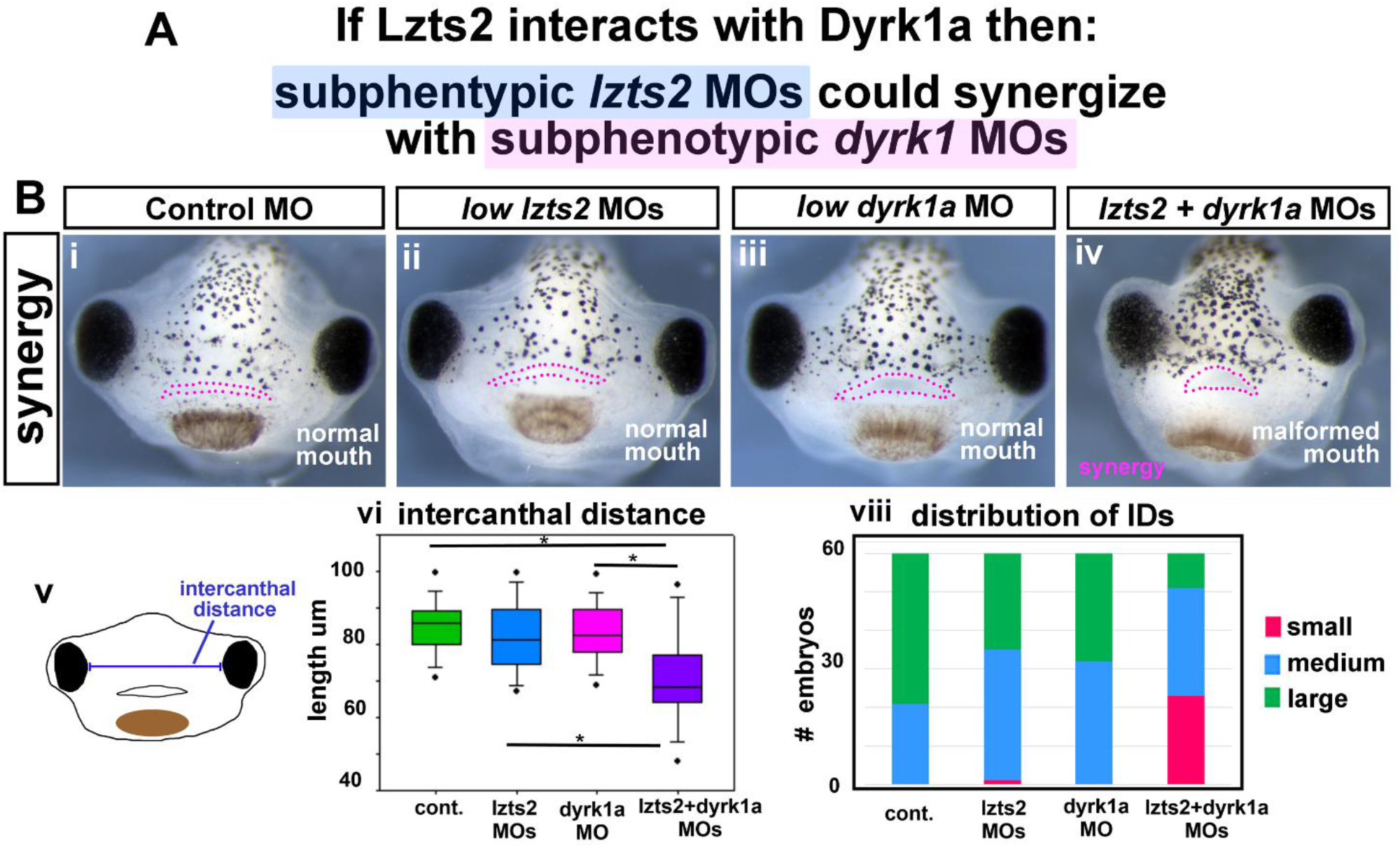
A) Sub-phenotypic reduction of Lzts2 and Dyrk1a synergistically disrupts craniofacial morphology. **B)** (i– iv) Frontal views of representative embryos injected with sub-phenotypic concentrations of control MOs, low-dose lzts2 MOs, low-dose dyrk1a MOs, or a combination of low-dose lzts2 and dyrk1a MOs. Individual injections produce normal craniofacial morphology, whereas combined knockdown results in malformed mouths and craniofacial defects. (v) Schematic illustrating measurement of intercanthal distance. (vi) Quantification of intercanthal distances for each condition (n = 60 embryos). (viii) Distribution analysis of intercanthal distances shown as stacked bar graphs categorized into small, medium, and large groups.

These results indicated that injection of a combination of *lzts2* and *dyrk1a* MOs resulted in phenotypes that are consistent with a synergistic effect supporting the possibility that Lzts2 and Dyrk1a could cooperate during craniofacial development.

### 6. Knockdown of Lzts2 can reduce the severity of craniofacial malformations in embryos with increased Dyrk1a

If LZTS2 is required for DYRK1A function, then reducing Lzts2 levels might be expected to attenuate craniofacial defects caused by excess DYRK1A (Fig. 5A). To test this possibility, we performed a phenotypic modifier assay in which embryos were injected with *dyrk1a* mRNA alone or in combination with Lzts2 MOs. As expected, control embryos injected with *GFP* mRNA and control MOs displayed normal craniofacial morphology (Fig. 5Bi). In contrast, embryos injected with dyrk1a mRNA (300 pg) together with control MOs (30–40 ng/embryo) developed prominent craniofacial abnormalities, including small eyes, a narrow midface, and malformed mouths (Fig. 5Bii). Embryos injected with Lzts2 MO (30–40 ng/embryo) and *GFP* mRNA (300 pg) displayed the moderate craniofacial phenotype described above. Although some malformations were shared between *dyrk1a*-overexpressing embryos and Lzts2 morphants, several features were clearly distinct. For example, the eyes of Lzts2 morphants were slanted, whereas embryos overexpressing *dyrk1a* exhibited reduced eye size. The shape of the mouth also differed between these conditions. Strikingly, embryos injected with both *dyrk1a* mRNA and Lzts2 MOs frequently displayed craniofacial defects that appeared less severe than those observed in embryos injected with either reagent alone (representative example Fig. 5Biv). While overall facial size remained reduced, the shape of the mouth appeared partially restored in a subset of embryos.

**Fig 5.**
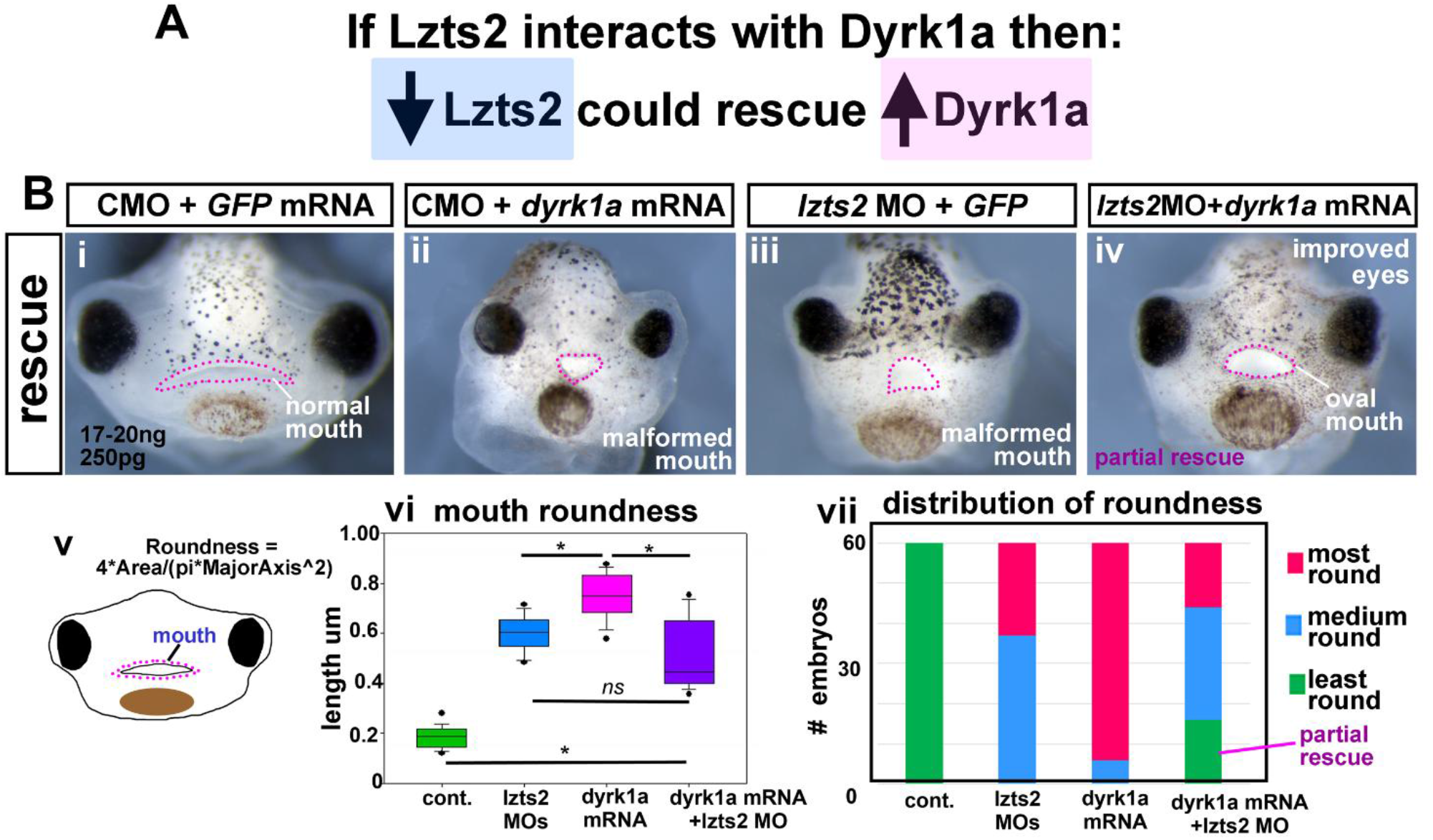
**A)** Reduction of Lzts2 partially suppresses craniofacial defects caused by Dyrk1a overexpression. **B)** (i–iv) Frontal views of representative embryos injected with control MOs + GFP mRNA, control MOs + dyrk1a mRNA, lzts2 MOs + GFP mRNA, or *lzts2* MOs + *dyrk1a* mRNA. Co-injection of lzts2 MOs with dyrk1a mRNA partially improves craniofacial morphology relative to embryos overexpressing dyrk1a alone. (v) Schematic illustrating measurement of mouth roundness.(vi) Quantification of mouth roundness for each condition (n = 60 embryos). (vii) Distribution of mouth roundness values presented as stacked bar graphs divided into most round, medium round, and least round categories.

To quantify these qualitative observations, we measured mouth roundness using ImageJ, where a value of 1 represents a perfect circle and values approaching 0 indicate a more elongated ellipse (Fig. 5Bv). Control embryos exhibited the expected elliptical mouth shape with an average roundness value of 0.186 (Fig. 5Bvi). Embryos injected with either *dyrk1a* mRNA or Lzts2 MO displayed altered mouth morphology, with increased roundness values of 0.748 and 0.598, respectively (Fig. 5Bvi). In embryos injected with both dyrk1a mRNA and Lzts2 MOs, the average roundness was 0.509 (Fig. 5Bvi). Although variability was high, reflecting improvement in only a subset of embryos, the average mouth roundness in the dyrk1a mRNA + Lzts2 MO group was significantly different from embryos injected with dyrk1a mRNA and control MOs (p < 0.001, Fig. 5Bvi). To further illustrate this effect, mouth roundness values were also analyzed as distributions, divided into most round, round, and least round categories and displayed as stacked bar graphs. Notably, the dyrk1a mRNA + Lzts2 MO group contained a subset of embryos within the least-round category, a class otherwise observed only in control embryos (Fig. 5Bvii). This distribution analysis supports our qualitative observations and indicates that reducing Lzts2 partially suppresses the craniofacial defects caused by excess DYRK1A in a subset of embryos.

Together, these results indicate that reducing Lzts2 can partially ameliorate the defects in mouth morphology caused by excess Dyrk1a in a subset of embryos. However, this suppression was not observed for all craniofacial features. For example, the intercanthal distance, which reflects the width of the midface, was not significantly altered in embryos injected with both lzts2 MOs and *dyrk1a* mRNA (nor shown). Several explanations could account for this incomplete rescue. One possibility is that technical limitations, such as variability in injection efficiency, reduce our ability to detect stronger modifier effects. Alternatively, the results may reflect the biological nature of Lzts2 regulation of Dyrk1a. Lzts2 may mediate Dyrk1a function only within specific tissues, during a restricted developmental window, or for a subset of Dyrk1a-dependent cellular processes during craniofacial morphogenesis. Consistent with this possibility, full suppression of the *dyrk1a* overexpression phenotype may require coordinated modulation of additional components of the Dyrk1a regulatory networks that operate during craniofacial development.

### 6. The Xenopus laevis Lzts2 protein is similar to the human LZTS2 protein

To provide some evidence that LZTS2 may have conserved function in humans the *Xenopus laevis* protein was compared to human LZTS2. Both of the *Xenopus laevis* Lzts2 homeologs (NCBI Accession #s XP_018080722.1, XP_041426885.1) shared 51% identities and 63% positives with human LZTS2 (NCBI Accession # AAH58938.1, Fig. 6). Importantly, an analysis of conserved domains identified 2 regions or domains conserved with humans (Fig. 6B). One such domain was a Fez1 domain which is the defining feature of the protein and that puts this protein in the Fezzin family (Fig. 6B). The second domain identified in both *Xenopus laevis* Lzts2 homeologs as well the human protein is a SMC_prok_A super family domain. This is reported to be a chromosome segregation ATPase which is involved in cell cycle control, cell division and chromosome partitioning (Fig. 6B). Using a Swiss Model (Expasy) we visually compared the predicted 3D structures of the human protein with the *Xenopus laevis* homeolog Lzts2.L. Results indicated very similar overall structure having almost identical alpha-helixes (Fig. 6C). Together, these results indicate that the *Xenopus laevis* Lzts2 proteins are similar to the human LZTS2 protein, suggesting the possibility of conserved function that will need experimental validation.

**Figure 6.**
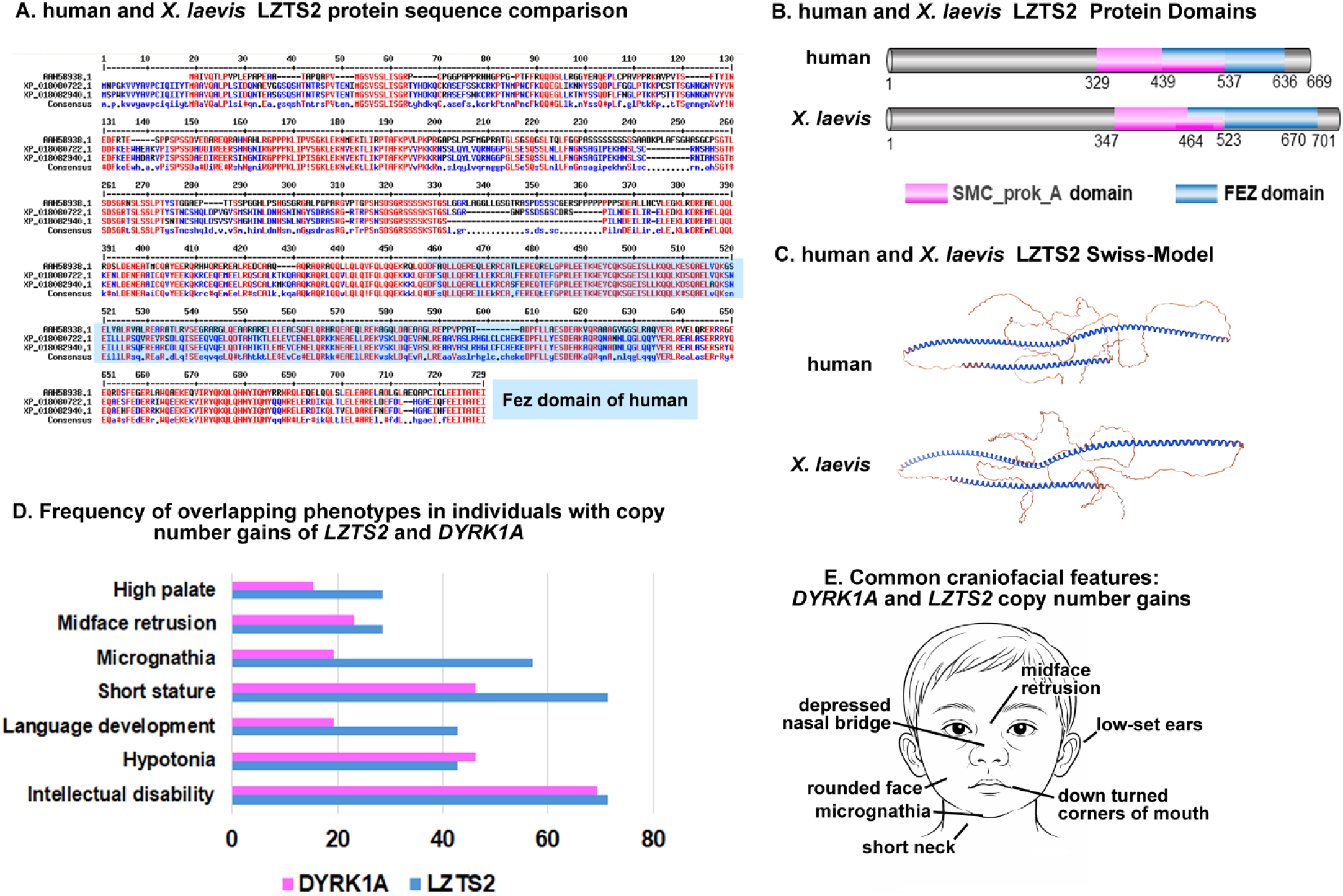
Evidence supporting a conserved function in humans. **A)** *Protein sequence comparison between human and Xenopus LZTS2*. Alignment of human LZTS2 (NCBI Accession #AAH58938.1) with the two *Xenopus laevis* Lzts2 homeologs (XP_018080722.1 and XP_041426885.1). The sequences show substantial conservation, with approximately 51% identity and 63% similarity (positives) between the *X. laevis* proteins and human LZTS2. Conserved residues are highlighted across the alignment. The region corresponding to the FEZ domain of the human protein is indicated. **B)** *Conserved protein domains in human and Xenopus LZTS2*. Schematic diagrams showing the domain organization of human LZTS2 and *X. laevis* Lzts2 proteins. Two conserved domains are present in both species: the SMC_prok_A domain (magenta), associated with chromosome segregation and cell-cycle related ATPase activity, and the FEZ domain (blue), a defining feature of Fezzin family proteins. The relative positions of these domains are similar between human and *Xenopus* proteins. **C)** Predicted structural comparison of human and Xenopus LZTS2. Predicted three-dimensional structures generated using Swiss-Model (Expasy) for human LZTS2 and the *X. laevis* Lzts2.L homeolog. The models display similar overall structural organization with comparable α-helical regions, indicating strong structural conservation between the species. **D)** Bar graph comparing the frequency of phenotypes (%) observed in individuals with LZTS2 copy number gains (blue, n=7) and DYRK1A copy number gains (magenta, n=26). Analysis was restricted to phenotypes present in two or more LZTS2 cases. Frequencies are expressed as the percentage of affected individuals within each cohort. **E)** Schematic showing common craniofacial phenotypes in individuals with *DYRK1A* copy number gains with one individual with copy number gain of *LZTS2*.

### 7. Phenotypic Overlap Between the LZTS2 and DYRK1A Gene Copy Number Gains

To explore whether LZTS2 and DYRK1A could possibly interact in human development we performed a comparative analysis of phenotypes associated with copy number gains in each gene to assess the extent of phenotypic convergence. Of 25 phenotypes identified in individuals with LZTS2 copy number gains, 24 (96%) were directly observed in at least one individual with DYRK1A copy number gain.

To further evaluate the strength of this overlap, we focused on phenotypes observed in two or more individuals with LZTS2 copy number gains and compared their frequencies to those reported in DYRK1A duplication cases (Fig. 6D). This analysis revealed that the most recurrent LZTS2-associated phenotypes align closely with high-frequency DYRK1A traits. Intellectual disability was the most prevalent feature in both cohorts followed by hypotonia and short stature. Craniofacial abnormalities also showed strong concordance, including micrognathia, midface retrusion, and high palate. Delayed language development was similarly enriched in both groups. These shared features cluster within neurodevelopmental and craniofacial domains. Interestingly, one individual with an LZTS2 copy number gain exhibited a particularly extensive constellation of craniofacial abnormalities that were also frequent in people with DYRK1A copy number gains and characteristic traits of Down Syndrome. Such traits included micrognathia, mid face retrusion, depressed nasal bridge, low-set ears, high or narrow palate, round facial morphology, and short neck and down turned corners of mouth (Fig. 6E).

Together, these findings demonstrate that LZTS2 copy number gains recapitulate the core phenotypic spectrum associated with DYRK1A dosage imbalance and are consistent with a potential functional interaction between these proteins during human development.

## DISCUSSION

Craniofacial development is governed by tightly regulated signaling networks in which dosage-sensitive factors play critical roles. While proteins such as DYRK1A have well-established functions in this process, less is known about the regulatory partners that modulate their activity during development. Our findings identify LZTS2 as both a contributor to craniofacial morphogenesis and a functional partner of DYRK1A, linking these proteins within a shared developmental framework that shapes craniofacial patterning.

### LZTS2 has a role in craniofacial development

Our findings demonstrate that LZTS2 is required for normal craniofacial development in Xenopus. Reduction of Lzts2 disrupted craniofacial morphogenesis, resulting in defects in eye shape, mouth morphology, and midfacial width. To date, in vivo studies of Lzts2 loss of function in mice remain limited, with a single developmental study demonstrating a critical role in kidney and urinary tract morphogenesis (Peng et al., 2011). Similarly, zebrafish studies are sparse, with one report showing that Lzts2 regulates gastrulation movements, dorsoventral patterning, and early organ progenitor specification through modulation of Wnt/β-catenin signaling (Li et al., 2011). Although craniofacial development was not explicitly examined in these models, published images reveal abnormalities in head and eye morphology (Li et al., 2011; Peng et al., 2011). Consistently, by examining published images and databases we found that LZTS2 is expressed in developing craniofacial tissues (Gessert et al., 2011; Li et al., 2011; Peng et al., 2011). Together, these findings support a previously underappreciated and potentially conserved role for LZTS2 in craniofacial development across vertebrate species.

### LZTS2 interacts with DYRK1A in craniofacial development

The craniofacial abnormalities observed in Lzts2 morphants closely resembled those resulting from reduced Dyrk1a, suggesting that these proteins may function within a shared developmental pathway. Supporting this idea, Lzts2 and Dyrk1a exhibited overlapping temporal and spatial expression during key stages of craniofacial development, consistent with the possibility that they act within the same cellular contexts in developing facial tissues. Functional interaction between these proteins was further supported by genetic interaction experiments. Sub-phenotypic reductions of Lzts2 and Dyrk1a synergized to produce pronounced craniofacial defects, indicating that partial disruption of both pathways is sufficient to impair normal morphogenesis. In contrast, partial reduction of Lzts2 attenuated aspects of the phenotype induced by Dyrk1a overexpression, suggesting that LZTS2 may modulate or constrain DYRK1A activity during development. Together, these findings support a functional interaction between LZTS2 and DYRK1A in craniofacial development and raise the possibility that LZTS2 acts as a context-dependent regulator of DYRK1A-mediated signaling during embryogenesis.

Interestingly, analysis of human copy number gain phenotypes provides independent support for this model. Individuals with duplications encompassing LZTS2 exhibit a constellation of neurodevelopmental and craniofacial features that closely mirror those associated with DYRK1A copy number gains, including intellectual disability, hypotonia, short stature, micrognathia, midface retrusion, and palate abnormalities. The observation that at least one individual with LZTS2 duplication presents with a combination of craniofacial features highly characteristic of DYRK1A-associated conditions as well as Down Syndrome further strengthens this relationship. While these human genetic data do not establish direct mechanistic interaction, their strong phenotypic concordance, when considered alongside the experimental evidence presented here, supports a model in which LZTS2 and DYRK1A function within a common developmental framework and that alterations in either gene can perturb overlapping pathways governing craniofacial development.

### Modulating DYRK1A through LZTS2 as a potential strategy for craniofacial disorders

Modulating DYRK1A activity has emerged as a potential therapeutic strategy for disorders associated with abnormal DYRK1A dosage. Indeed, reducing DYRK1A activity has shown promise in several preclinical models of Down syndrome (Becker et al., 2014; De Toma et al., 2019; Feki and Hibaoui, 2018; Neumann et al., 2018; Nguyen et al., 2018; Souchet et al., 2015; Stotani et al., 2016; Stringer et al., 2017). For example, genetic or pharmacological normalization of Dyrk1a levels can partially improve craniofacial abnormalities in mouse models of Down syndrome (McElyea et al., 2016; Redhead et al., 2023). However, broad inhibition of DYRK1A presents important challenges. In mouse models of Down syndrome, altering DYRK1A activity can itself disrupt craniofacial development depending on the dose and developmental timing of inhibition (Jamal et al., 2022; Starbuck et al., 2021). In addition, currently available DYRK1A inhibitors may lack complete specificity and can produce off-target effects, while pharmacokinetic factors such as tissue diffusion and metabolism may further complicate therapeutic outcomes (Stringer et al., 2017). Because DYRK1A participates in numerous cellular processes, including functions associated with tumor suppression, broad inhibition of the kinase could have unintended consequences(Ananthapadmanabhan et al., 2023).

An alternative strategy may therefore be to modulate regulatory proteins that influence DYRK1A activity or localization, rather than directly inhibiting the kinase itself. Increasing evidence suggests that the spatiotemporal regulation of kinase signaling, including that of DYRK1A, is strongly influenced by scaffolding and adaptor proteins that organize regulatory complexes (Huber and Brummer, 2024). In the present study, we provide experimental evidence that Lzts2 can modify Dyrk1a function during craniofacial development. Using *Xenopus laevis* embryos, we found that reducing Lzts2 in embryos overexpressing *dyrk1a* resulted in a partial improvement in mouth morphology, suggesting that modulation of Lzts2 can influence the phenotypic consequences of excess DYRK1A. Importantly, this improvement was limited to a subset of embryos and specific craniofacial features. These observations raise the possibility that Lzts2 regulates only particular cellular functions or developmental contexts in which Dyrk1a operates. If so, targeting Lzts2 or related regulatory components could provide a means to selectively modify specific outputs of DYRK1A signaling, rather than broadly suppressing all DYRK1A activity. Such an approach may allow fine-tuning of DYRK1A-dependent pathways while preserving its many essential cellular functions. In addition, modulation of other components of DYRK1A regulatory complexes could further refine distinct aspects of DYRK1A signaling, providing a framework in which different interacting proteins selectively tune specific developmental outputs of the kinase.

### Limitations of the study and future directions

Several limitations of this study should be considered when interpreting these findings. First, although the *Xenopus laevis* embryo provides a powerful system for studying craniofacial development and genetic interactions in vivo, developmental mechanisms identified in amphibian embryos may not fully recapitulate those operating in mammalian craniofacial tissues. While we demonstrate that the Xenopus Lzts2 protein shares strong sequence and structural similarity with human LZTS2, additional studies in mammalian systems will be important to determine whether similar regulatory interactions occur during human craniofacial development. A key limitation in the phenotypic analysis of humans is that copy number gains represent duplications of genomic segments that often encompass multiple genes in addition to LZTS2, making it difficult to attribute specific phenotypes solely to LZTS2 dosage. Furthermore, variability in duplication size, genetic background, and clinical reporting across individuals may contribute to phenotypic heterogeneity and limit precise genotype–phenotype correlations. Further biochemical and cell biological studies will be required to determine whether Lzts2 directly regulates Dyrk1a localization, stability, or interaction with other regulatory partners and additional components of Dyrk1a signaling complexes. Finally, because LZTS2 has been shown to regulate β-catenin and Wnt signaling in other cellular contexts (Thyssen et al., 2006), it remains possible that the effects observed in this study reflect indirect regulation of Dyrk1a through changes in Wnt-dependent signaling pathways. Future work will therefore be required to distinguish between direct interaction of Dyrk1a and Lzts2 and indirect effects mediated through broader signaling networks that influence craniofacial development.

## Conclusion

This work identifies Lzts2 as an essential regulator of craniofacial development and demonstrates that it functionally interacts with Dyrk1a during formation of this region. These findings suggest that Lzts2 contributes to the Dyrk1a-dependent signaling in a developmental context. Understanding how regulatory proteins such as Lzts2 shape Dyrk1a activity may provide new opportunities to selectively tune specific DYRK1A-dependent pathways, enabling more precise therapeutic strategies for disorders, such as Down Syndrome, associated with abnormal DYRK1A dosage.

## Acknowledgements

We thank our animal care technician, Leona Bhandari, for her essential support. We are also grateful to the VCU Children’s Hospital Research Institute for funding that made this work possible and fostered this collaboration.

## FIGURES AND LEGENDS

**Supplementary Figure 1.**
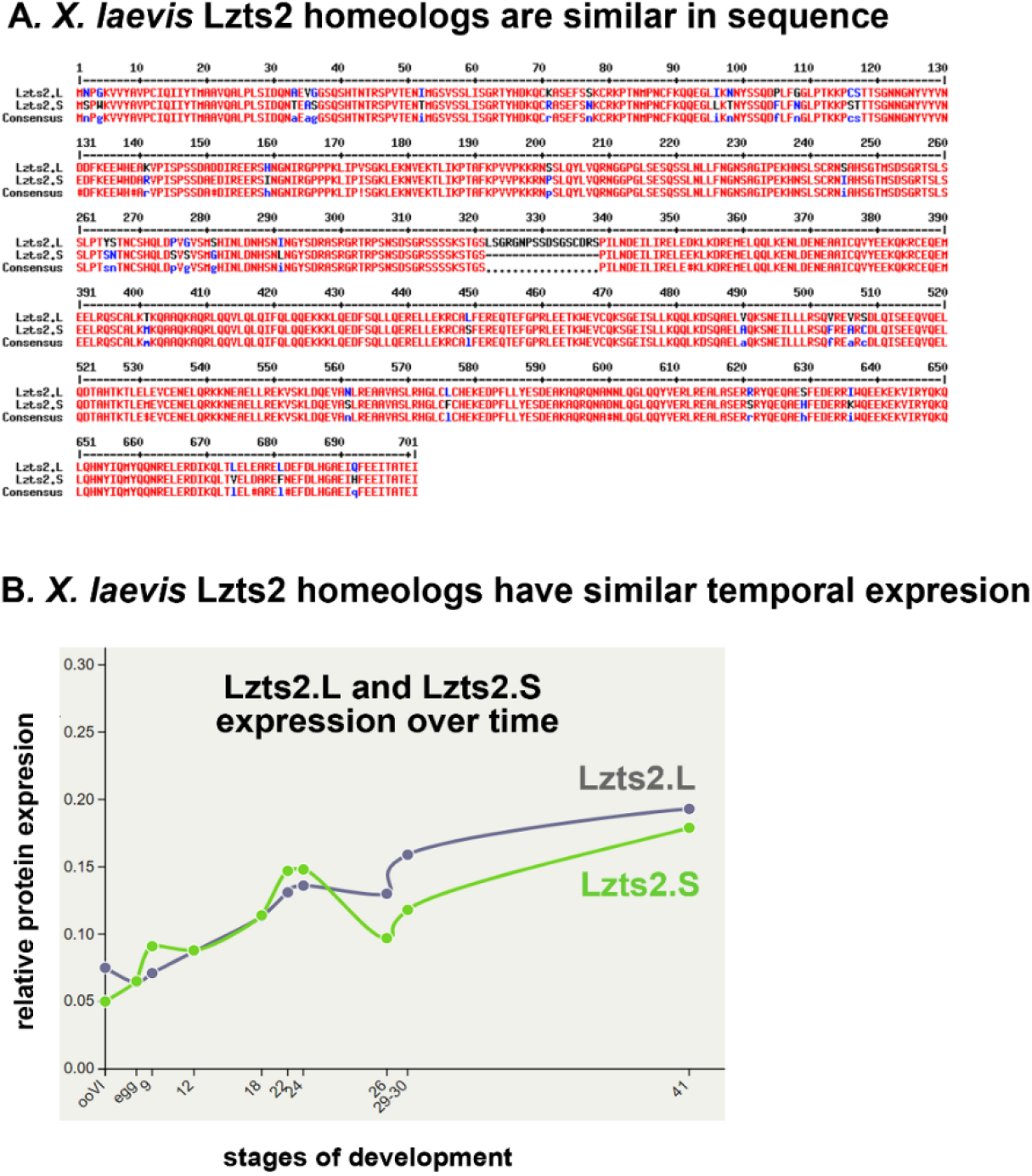
Xenopus laevis Lzts2 homeologs are highly similar in sequence and expression. **A)** *Sequence alignment of Xenopus laevis lzts2 homeologs*. Protein sequence alignment comparing the two *X. laevis lzts2* homeologs (*lzts2*.*L* and *lzts2*.*S*). Conserved residues are highlighted, demonstrating a high degree of sequence similarity between the two proteins. **B)** *Temporal expression of Lzts2 homeologs during development*. Relative protein expression levels of Lzts2.L and Lzts2.S across developmental stages in *X. laevis*. Both homeologs display similar temporal expression profiles throughout embryogenesis.

